# Restoration of wild-type motility to flagellin-knockout *Escherichia coli*

**DOI:** 10.1101/700492

**Authors:** Nicholas M. Thomson, Mark J. Pallen

## Abstract

Flagellin is the major constituent of the flagellar filament and faithful restoration of wild-type motility to flagellin mutants may be beneficial for studies of flagellar biology and biotechnological exploitation of the flagellar system. Therefore, we explored the restoration of motility by flagellin expressed from a variety of combinations of promoter, plasmid copy number and induction strength. Motility was only partially restored using the tightly regulated rhamnose promoter, but wild-type motility was achieved with the T5 promoter, which, although leaky, allowed titration of induction strength. Motility was little affected by plasmid copy number when dependent on inducible promoters. However, plasmid copy number was important when expression was controlled by the native *E. coli* flagellin promoter. Motility was poorly correlated with flagellin transcription levels, but strongly correlated with the amount of flagellin associated with the flagellar filament, suggesting that excess monomers are either not exported or not assembled into filaments. This study provides a useful reference for further studies of flagellar function and a simple blueprint for similar studies with other proteins.

## 1. Introduction

Restoration of gene function in a knockout strain by *trans*-complementation is widely used to link genes to phenotypes. Similarly, expression of an engineered variant of a protein from a plasmid is widely used for biotechnological purposes. However, plasmid-based expression can disassociate a gene from its wild-type promoter and other regulatory influences and can lead to changes in gene copy number; all of which may lead to unwanted changes in gene expression and phenotype. This is a particular problem in synthetic biology, which relies on well-characterised, predictable gene expression when multiple genetic parts are joined to create new pathways, networks and products.

The flagellar filament in *Escherichia coli* is primarily responsible for swimming motility, but has also been implicated in interactions with host cells and the immune system [1,2]. The filament is composed of multiple subunits of a single protein, flagellin (FliC), which travel in a semi-folded state through the hollow filament to assemble at the distal tip [3]. Interest has focused on engineering flagellin to display protein domains in the solvent-exposed regions [4,5], which typically requires wild-type levels of expression of the flagellin gene from plasmids. However, when this has been attempted in previous studies, flagellar motility in the complemented strain has either not been compared to that in an isogenic wild-type strain [6–8], or where they have been compared, the complemented strain shows reduced motility [9]. We therefore sought to establish a baseline for effective restoration of wild-type motility in *fliC*-deficient *E. coli* by comparing expression of flagellin from a variety of promoters and plasmids. Although our investigation focussed on motility, the general approach is easy to adopt and strongly recommended for any plasmid-based complementation system to ensure confident and reliable interpretation of experimental results.

## 2. Material and methods

### 2.1. Bacterial strains, plasmids, and growth conditions

The bacterial strains and plasmids used in this study are summarised in Table 1. *E. coli* strain AW405 was a gift from Howard Berg (Harvard University). AW405 is resistant to streptomycin due to a *rpsL136* chromosomal mutation. Therefore, streptomycin (100 mg.mL^−1^) was included in all AW405 and AW405Δ*fliC* cultures as a precaution against contamination. Antibiotics were also added at the following concentrations as required for plasmid maintenance and selection (Table 1): carbenicillin (100 mg.mL^−1^), kanamycin (50 mg.mL^−1^), chloramphenicol (30 mg.mL^−1^). Unless otherwise specified, cultures were grown in 5 mL Miller’s Luria-Bertani (LB) medium in 30 mL polystyrene, screw-capped universal tubes, at 37 °C and shaking at 200 rpm. When growth on solid medium was required, agar (1.5%) was added to the LB medium and plates were incubated statically at 37 °C.

**Table 1:**
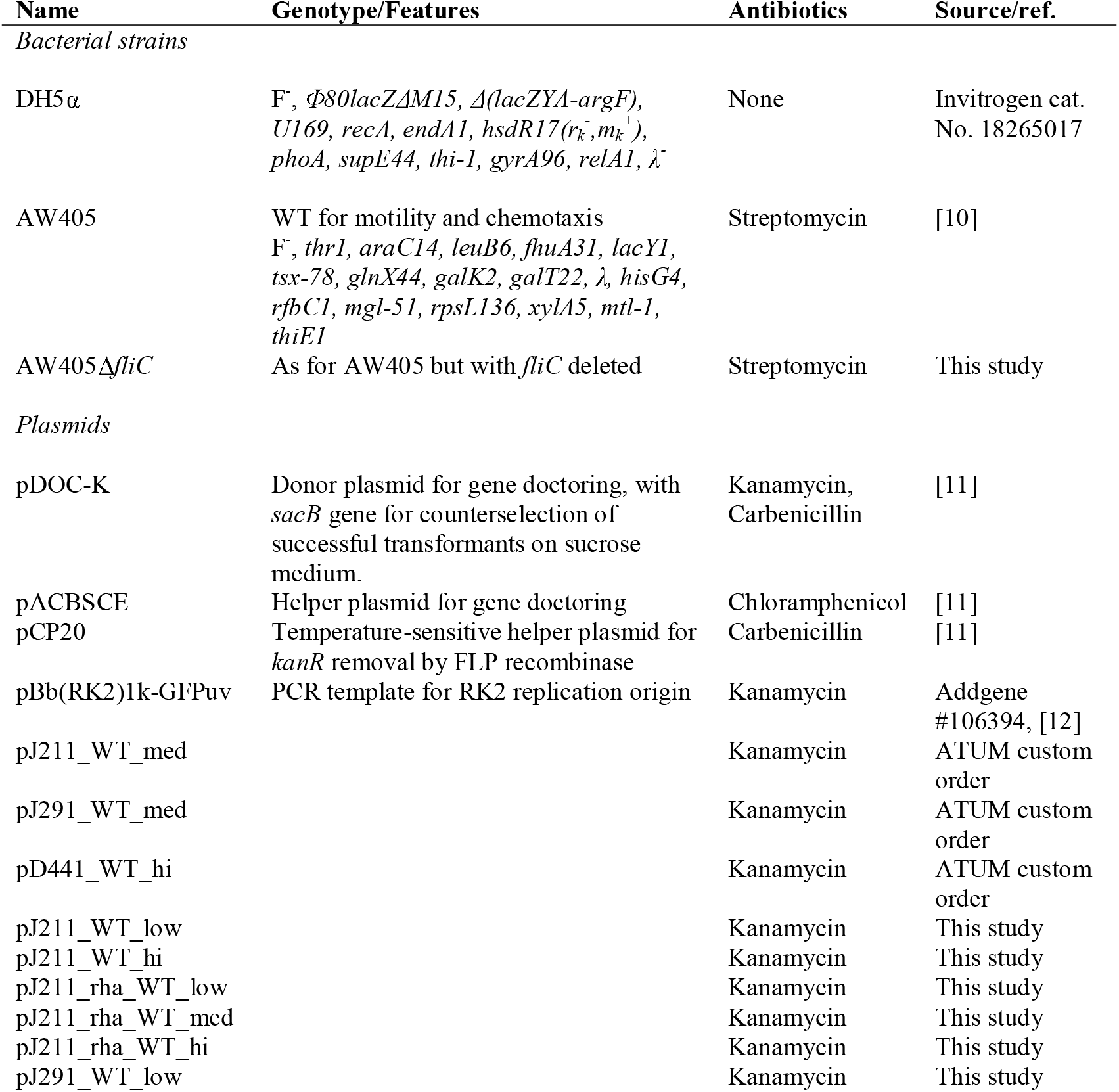
Bacterial strains and plasmids used in this study

| Name | Genotype/Features | Antibiotics | Source/ref. |
| --- | --- | --- | --- |
| *Bacterial strains* |  |  |  |
| DH5α | F, Φ80*lacZ*ΔM15, $\Delta$(*lacZYA-argF*), *U169*, *recA*, *endA1*, *hsdR17*($r_k^-$, $m_k^+$), *phoA*, *supE44*, *thi-1*, *gyrA96*, *relA1*, $\lambda^-$ | None | Invitrogen cat. No. 18265017 |
| AW405 | WT for motility and chemotaxis<br>F, *thr1*, *araC14*, *leuB6*, *fhuA31*, *lacY1*, *tsx-78*, *glnX44*, *galK2*, *galT22*, $\lambda$, *hisG4*, *rfbC1*, *mgl-51*, *rpsL136*, *xylA5*, *mtl-1*, *thiE1* | Streptomycin | [10] |
| AW405Δ*fliC* | As for AW405 but with *fliC* deleted | Streptomycin | This study |
| *Plasmids* |  |  |  |
| pDOC-K | Donor plasmid for gene doctoring, with *sacB* gene for counterselection of successful transformants on sucrose medium. | Kanamycin, Carbenicillin | [11] |
| pACBSCE | Helper plasmid for gene doctoring | Chloramphenicol | [11] |
| pCP20 | Temperature-sensitive helper plasmid for *kanR* removal by FLP recombinase | Carbenicillin | [11] |
| pBb(RK2)1k-GFPuv | PCR template for RK2 replication origin | Kanamycin | Addgene #106394, [12] |
| pJ211_WT_med |  | Kanamycin | ATUM custom order |
| pJ291_WT_med |  | Kanamycin | ATUM custom order |
| pD441_WT_hi |  | Kanamycin | ATUM custom order |
| pJ211_WT_low |  | Kanamycin | This study |
| pJ211_WT_hi |  | Kanamycin | This study |
| pJ211_rha_WT_low |  | Kanamycin | This study |
| pJ211_rha_WT_med |  | Kanamycin | This study |
| pJ211_rha_WT_hi |  | Kanamycin | This study |
| pJ291_WT_low |  | Kanamycin | This study |

### 2.2. Selecting combinations of promoters and replication origins

The expression plasmids were based on pre-existing laboratory stocks of pJ211_WT_med, pJ291_WT_med, and pD441_WT_hi, all of which were custom-made by ATUM (Newark, CA, U.S.A.). Each of the starting plasmids contained a copy of *fliC* from *E. coli* AW405 under control of either its natural promoter (pJ211) or the IPTG-inducible T5 promoter (pJ291 and pD441), and with either the medium copy-number p15a replication origin (med) or the high copy-number pUC origin (hi). To most closely replicate the low gene dosage of chromosome-encoded genes, we also selected the RK2 replication origin, which has previously been shown to consistently maintain plasmids with low copy numbers of ~5 per cell [13,14]. The two promoters on the set of starting plasmids provided for either endogenously-controlled or inducible expression. However, since *lac*-based promoters including T5 are not truly tuneable [15], we also chose to include the rhamnose promoter to investigate whether tuneability of expression levels on the individual cell level would have a different effect on motility to the all-or-nothing response of the T5 promoter. We therefore constructed a set of nine plasmids with all combinations of three promoters and three replication origins.

### 2.3. Plasmid construction

All cloning operations were performed using *E. coli* DH5α as the host strain. Plasmids were purified when necessary using a NucleoSpin Plasmid Miniprep kit (Macherey-Nagel, Düren, Germany). In the first stage of cloning, the *fliC* promoter was replaced with the rhamnose promoter in pJ211_WT_med. The plasmid was digested with SalI and NcoI and the linearised backbone was recovered by agarose gel purification. A gBlock containing the rhamnose promoter (from BioBrick BBa_K914003) and *E. coli rhaBAD* ribosome binding site together with SalI and NcoI restriction sites (Table 2) was purchased (IDT, Leuven, Belgium), digested and ligated to the backbone. This yielded pJ211_rha_WT_med.

**Table 2:**
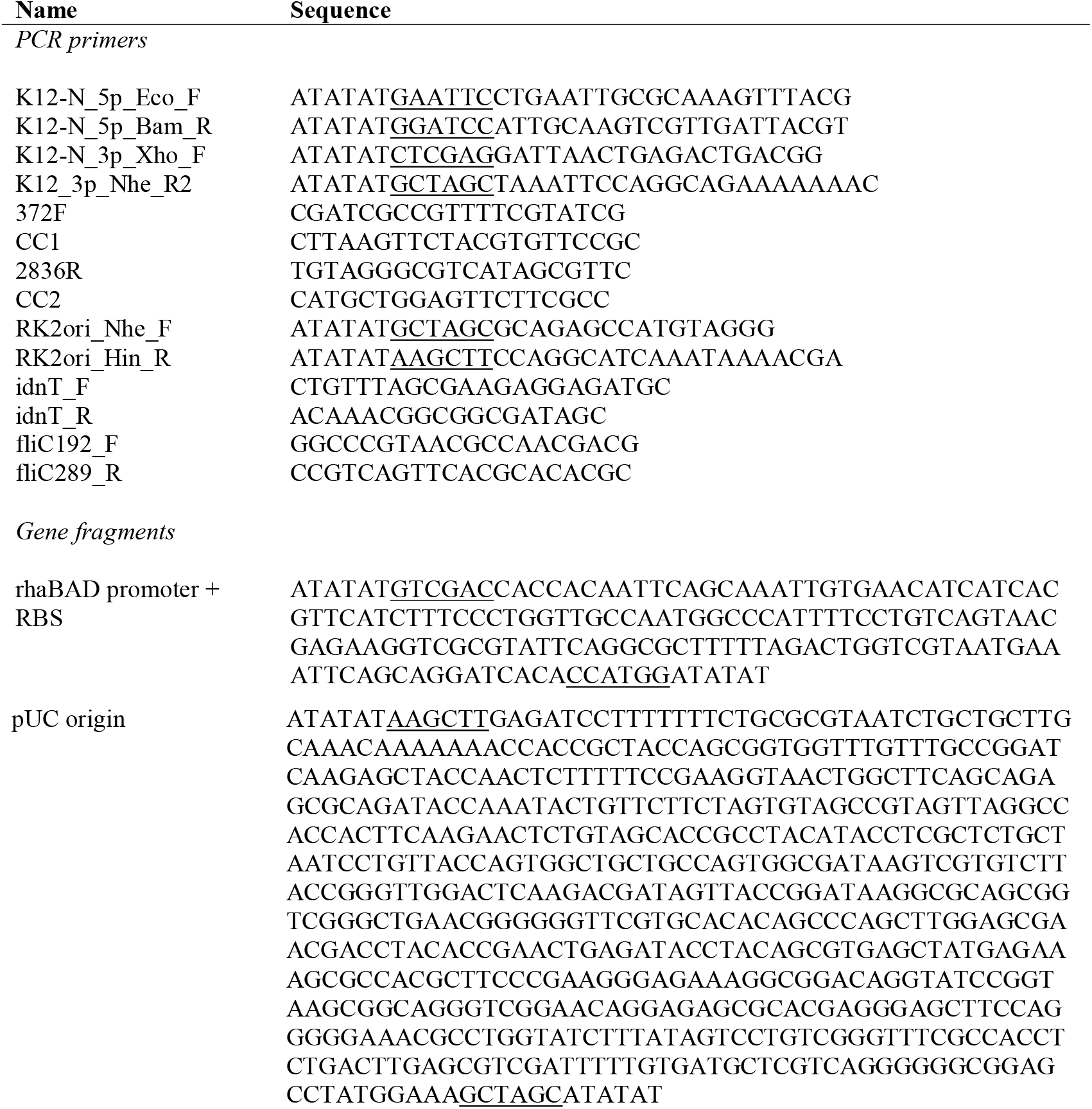
PCR primers and gene fragments used in this study. Underlined sequences indicate restriction sites.

In the second stage, the p15a replication origin was replaced by RK2 in each of the three medium copy plasmid variants. RK2 was amplified from pBb(RK2)1k-GFPuv [12], which was a gift from Brian Pfleger (University of Wisconsin-Madison), using Q5 DNA polymerase and primers RK2ori_Nhe_F and RK2ori_Hin_R (Table 2). The product was digested with NheI and HindIII, gel purified and ligated into the gel purified plasmid backbones, which had been digested with the same enzymes. This yielded pJ211_WT_low, pJ211_rha_WT_low and pJ291_WT_low.

Finally, pJ211_WT_med and pJ211_rha_WT_med were used for replacement of the p15a replication origin by the pUC origin. This proceeded as for insertion of RK2, except the pUC origin sequence was purchased as a gBlock (IDT,Leuven, Belgium) containing NheI and HindIII restriction sites (Table 2). The resulting plasmids were pJ211_WT_hi and pJ211_rha_WT_hi. All nine plasmid sequences were verified by Illumina shotgun sequencing in the DH5α host cells and mapping reads to the expected plasmid sequence. The function of the promoter-flagellin combinations were verified by performing a motility assay on cells of AW405Δ*fliC* transformed with each plasmid. Motile cells from each assay were recovered and grown overnight to produce glycerol stocks, which were used for all other experiments.

### 2.4. Creation of a fliC knockout strain

A *fliC* deletion mutant of AW405 was created using the gene doctoring system for chromosomal gene editing [11]. Plasmids pDOC-K, pACBSCE and pCP20 were gifts from Stephen Busby (University of Birmingham). Briefly, homologous regions of 200 bp upstream and downstream of the *fliC* gene were cloned into pDOC-K using EcoRI/BamHI (upstream) and XhoI/NheI (downstream) restriction sites. The inserted sequences were acquired from colony PCR reactions with AW405 as the template, Q5 DNA polymerase (NEB, Hitchin, U.K.) and primers K12-N_5p_Eco_F, K12-N_5p_Bam_R (upstream), K12-N_3p_Xho_F, and K12_3p_Nhe_R2 (downstream) (Table 2), followed by agarose gel purification using a NucleoSpin Gel and PCR Cleanup kit (Macherey-Nagel, Düren, Germany).

The plasmids pDOC-K (containing the homologous regions) and pACBSCE were then inserted into AW405 by co-electroporation and cells containing both plasmids were selected for on LB agar containing kanamycin, carbenicillin and chloramphenicol. A single colony was picked, grown to late exponential phase in LB + antibiotics (0.5 mL), washed free of antibiotics in 0.1× LB and incubated at 37 °C for 1 h in the same volume of 0.1× LB containing 0.5% L-arabinose to induce expression of the λ-Red recombinase. Successful recombinants were selected for by plating on LB agar with 5% sucrose and screened for loss of both plasmids by checking susceptibility to carbenicillin and chloramphenicol. Deletion of the *fliC* gene was confirmed by colony PCR using OneTaq DNA polymerase (NEB, Hitchin, U.K.) and primers 372F, CC1 (upstream fragment), 2836R and CC2 (downstream fragment).

The *kanR* resistance cassette was removed by transformation with pCP20 and growth of carbenicillin resistant colonies in LB at 30 °C overnight. The temperature-sensitive pCP20 was then removed by growth at 37 °C without antibiotics. Colonies were finally checked for susceptibility to kanamycin, carbenicillin and chloramphenicol, and verified by colony PCR with primers 372F and 2836R, and shotgun sequencing.

### 2.5. Plasmid and genome sequence analysis

Genomic DNA was purified using a FastDNA Spin Kit for feces (MP Bio, Santa Ana, CA, U.S.A.) according to the manufacturer’s instructions, except that the final elution was in 200 μL of DNAse-free water rather than 60 μL. Plasmids were purified with a NucleoSpin Plasmid Miniprep kit. The DNA was quantified using a Quant-iT dsDNA high sensitivity assay kit (Thermo Fisher, Waltham, MA, U.S.A.) and normalised to 0.2 ng.μL^−1^ in 10 mM Tris-HCl. Sequencing libraries were prepared with the Nextera XT DNA Library Prep kit (Illumina, San Diego, CA, U.S.A.). Libraries were quantified using the Quant-iT dsDNA high sensitivity assay kit. Genome samples were pooled in equal quantities, while plasmid samples were first pooled together and the whole plasmid pool was added to the genomic pool at one-tenth the quantity of a genomic sample. The final pool was then run at a final concentration of 1.8 pM on an Illumina NextSeq 500 instrument using a mid-output sequencing kit for 150 bp paired-end reads. Sequences were quality checked by FastQC v0.11.7 and trimmed with Trimmomatic v0.36 with a minimum read length of 40 bp and a sliding window of 4 bp with average quality of 15. The reads were then mapped against the expected sequence for each sample in Geneious R11.1 with the Geneious mapper and default settings.

### 2.6. Motility assays

Motility assays were performed on motility agar (MA) plates, which consisted of (per L) Difco Lab-Lemco powder (10 g), Difco meat extract (3 g), NaCl (5 g) and agar (3 g). 30 mL of medium was solidified in a 90 mm triple-vented petri dish for each measurement. Kanamycin, L-rhamnose (Sigma-Aldrich, Gillingham, U.K.) and Isopropyl β-D-1-thiogalactopyranoside (IPTG; Sigma-Aldrich, Gillingham, U.K.) were included at the specified concentrations whenever necessary. A master plate of each strain was initially prepared by stabbing from a glycerol stock into the centre of a MA plate and incubating at 30 °C overnight. Subsequent inoculations were made from the outer edge of the motile disk on the master plate.

Three biological replicates were grown for each strain and three technical replicates were inoculated from each culture. The cultures were grown in motility broth (MB; as for MA but omitting the agar) at 30 °C overnight, then normalised to OD_600_ = 0.5. 1 μL of normalised culture was applied to the centre of a MA plate by stabbing a 10 μL pipette tip into the agar, taking care not to reach through to the bottom, and ejecting the inoculum. Plates were then incubated at 30 °C for 24 h and motility was quantified by measuring the diameter of the motile disk. All assays from each promoter type were performed using the same batch of MA, and each biological replicate was performed on the same day. Due to the volume of media required it was not possible to perform all assays simultaneously, and this gave rise to some batch-to-batch variation. Therefore, an AW405 wild-type control was performed alongside each set of plates, and the final motility scores were scaled according to the mean of all wild-type assays (28.88 mm) to allow accurate comparisons between promoters.

### 2.7. Reverse transcriptase-quantitative polymerase chain reaction

Flagellin may be present in the cytoplasm (either as soluble or insoluble protein), assembled into flagellar filaments, or diluted in the external medium as unpolymerised monomers and fragments of broken filaments. Therefore, obtaining an accurate measure of total flagellin protein is challenging. However, promoter induction strength and plasmid copy number are expected to modulate flagellin expression at the transcriptional level, so we measured the levels of *fliC* transcription relative to the wild-type strain for each plasmid construct using reverse transcriptase-quantitative polymerase chain reaction (RT-qPCR). We attempted to recover RNA directly from the motile cultures on plates but a combination of relatively low available cell numbers and agar inhibiting cell lysis meant that we were unable to recover sufficient RNA. Therefore, we extracted RNA from liquid cultures in MB medium.

Starter cultures (in triplicate) were grown as for motility assays and then normalised to OD_600_ = 0.1. The normalised cultures were further diluted 10-fold in MB, then 100 μL was used to inoculate 10 mL of MB + streptomycin, kanamycin, L-rhamnose and IPTG as necessary and the cultures were grown for 15 h at 30 °C, 200 rpm. For the inducible promoters, we included the concentration of inducer that gave the greatest motility as reported in section 3.1, i.e. 0.5% rhamnose for the *rha* promoter and 100 μM IPTG for the T5 promoter. Positive and negative control cultures consisting of AW405 and AW405Δ*fliC*, respectively, were grown in the same way. RNA was recovered from the cultures using a RNeasy Power Microbiome RNA purification kit (Qiagen, Venlo, Netherlands). The RNA concentration was determined by Qubit RNA high sensitivity assay (Invitrogen, Inchinnan, U.K.) and integrity checked by TapeStation RNA assay (Agilent, Stockport, U.K.), then adjusted to 3.125 ng.μL^−1^ with nuclease-free water.

Primers were designed in Geneious Prime 2019. *fliC* primers were fliC_192_F and fliC_289_R (Table 2). The endogenous control was *idnT* [16] and used primers idnT_F and idnT_R (Table 2). Reactions were set up in 384-well microplates (4titude, Dorking, U.K.) and consisted of 2.5 μL PrecisionPLUS OneStep RT-qPCR Master Mix with ROX and SYBR green (Primer Design, Chandler’s Ford, U.K.), 0.3 μM final concentration of each primer, 2 μL normalised RNA solution and water to a final volume of 5 μL. Calibration curves for each target were constructed by triplicate reactions of seven 5-fold serial dilutions of purified AW405 gDNA (starting from 40 ng.μL^−1^). Reactions were run with 3 technical replicates each on a ViiA7 Real-time PCR system (Thermo Fisher, Waltham, MA, U.S.A.) programmed as follows: 55 °C for 10 min, 95 °C for 2 min, 50 cycles of 95 °C for 10 sec and 60 °C for 60 sec, then a melt curve analysis using 95 °C for 15 sec, 60 °C for 60 sec and ramping at 0.05 °C.sec^−1^ to 95 °C. Data were analysed in QuantStudio v.1.3 (Thermo Fisher, Waltham, MA, U.S.A.) using the relative standard curve method.

### 2.8. SDS-polyacrylamide gel electrophoresis and densitometry

Triplicate cultures, including positive and negative controls, were grown in triplicate as described for RT-qPCR and the OD_600_ were measured. The cells were pelleted by centrifugation at 3000×g for 10 min. The culture with the lowest OD_600_ was resuspended in 40 μL phosphate buffered saline and the other cultures were resuspended in a volume to give an equal cell density. Flagella were depolymerised at 65 °C for 5 min in a water bath and the cells were removed by centrifugation as before. 25 μL of each supernatant was recovered, mixed with an equal volume of 2× Novex Tris-Glycine SDS Sample Buffer (Invitrogen, Inchinnan, U.K.) and 40 μL were run on an 8% Novex tris-glycine SDS-PAGE gel for 50 min at 200 V and stained using Bio-Safe Coomassie Stain (Bio-Rad, Watford, U.K.). The gels were photographed and the density of FliC bands (51.3 kDa) compared using the gel analyser function of ImageJ v1.52n. The 58 kDa band from the Color Prestained Protein Standard, Broad Range (NEB, Hitchin, U.K.) was used to normalise for staining differences between gels.

## 3. Results

### 3.1. Motility assays

We transformed each of the nine variant plasmids into *E. coli* AW405Δ*fliC* and conducted motility assays in 0.3% motility agar plates. The rhamnose promoter plasmids showed no motility in the absence of rhamnose (Figure 1 (A)). Motility increased in proportion to rhamnose concentration up to approximately 0.01% with a small further increase for higher concentrations, and reached a plateau at approximately 0.1%. The low copy number plasmid (RK2 origin) consistently gave the lowest motility, followed by the medium copy plasmid (p15a origin) and the high copy number plasmid (pUC origin) gave the highest motilities. However, the differences between plasmids were relatively small. In all cases, motility was significantly lower than for the WT control (T-test; p < 10^−4^).

**Figure 1:**
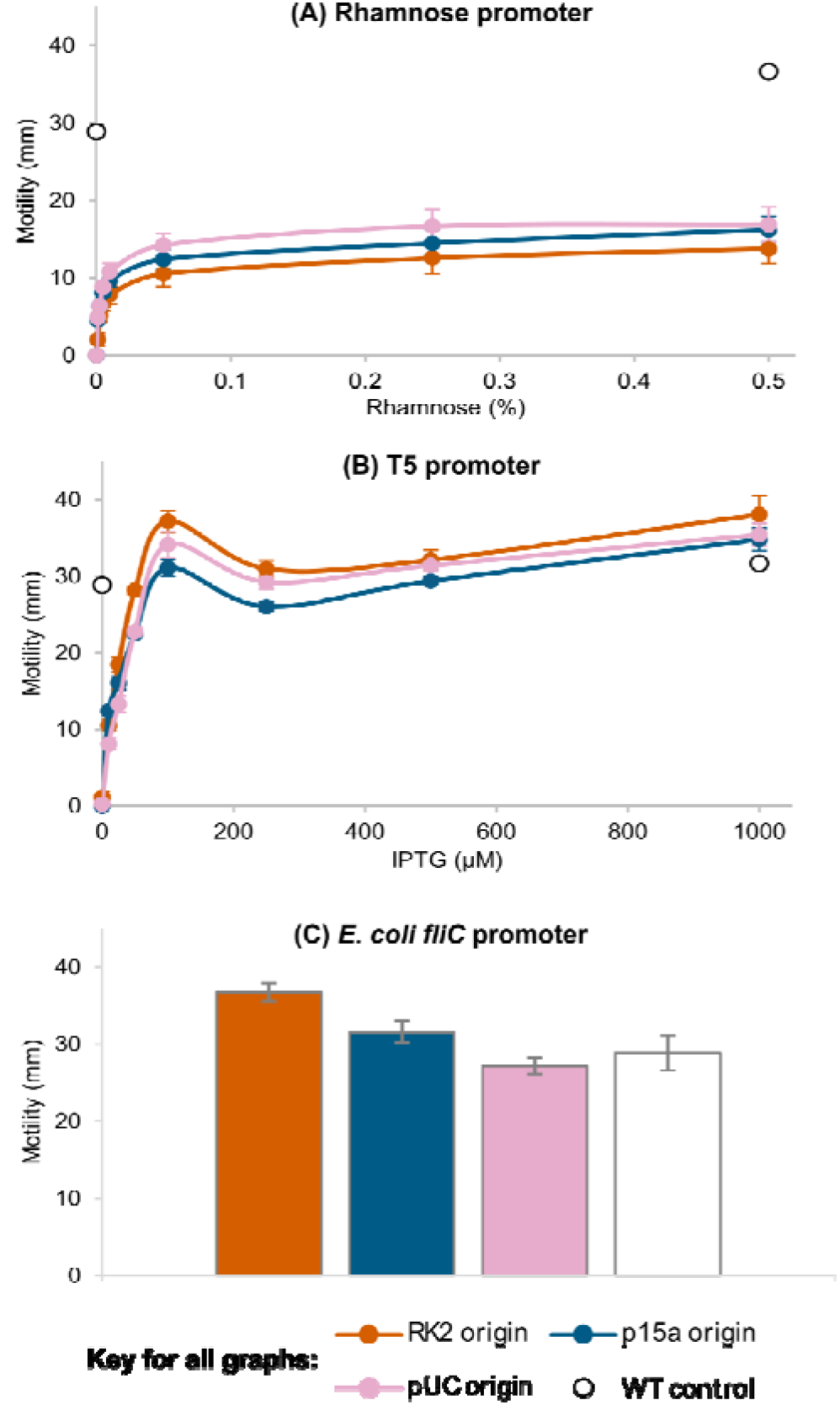
Variation in motility for expression of FliC from low (RK2), medium (p15a) and high (pUC) copy number plasmids with different combinations of promoter and induction strength. Motility was assayed by incubating inoculations for 24 h at 30 °C on soft motility agar plates and measured as the diameter of the motile swarm. (A) FliC under control of the rhamnose promoter, induced by a range of rhamnose concentrations. (B) FliC under control of the T5 promoter, induced by a range of IPTG concentrations. (C) FliC expressed from its natural promoter, which is not chemically induced. Wild-type strain AW405, with FliC expressed from the chromosome was assayed as a control. Results are the means of at least 3 biological repeats, each with 3 technical repeats ± SEM.

The T5 promoter also gave reasonably tight control over motility, although in a few cases leaky expression led to low levels of motility in the absence of IPTG. Motility increased with IPTG concentration up to approximately 100 μM (Figure 1 (B)). Interestingly, for each plasmid, 100 μM IPTG resulted in a local maximum motility, with lower motility at 250 mM. The motility then gradually increased again up to 1000 μM IPTG. The differences in motility between plasmids were small, with a consistent trend of p15a<pUC<RK2. Motilities peaked at above the level of the control, especially with the low copy number plasmid. However, for each plasmid it was possible to replicate wild-type motility by adjusting the concentration of IPTG (Figure 2).

**Figure 2:**
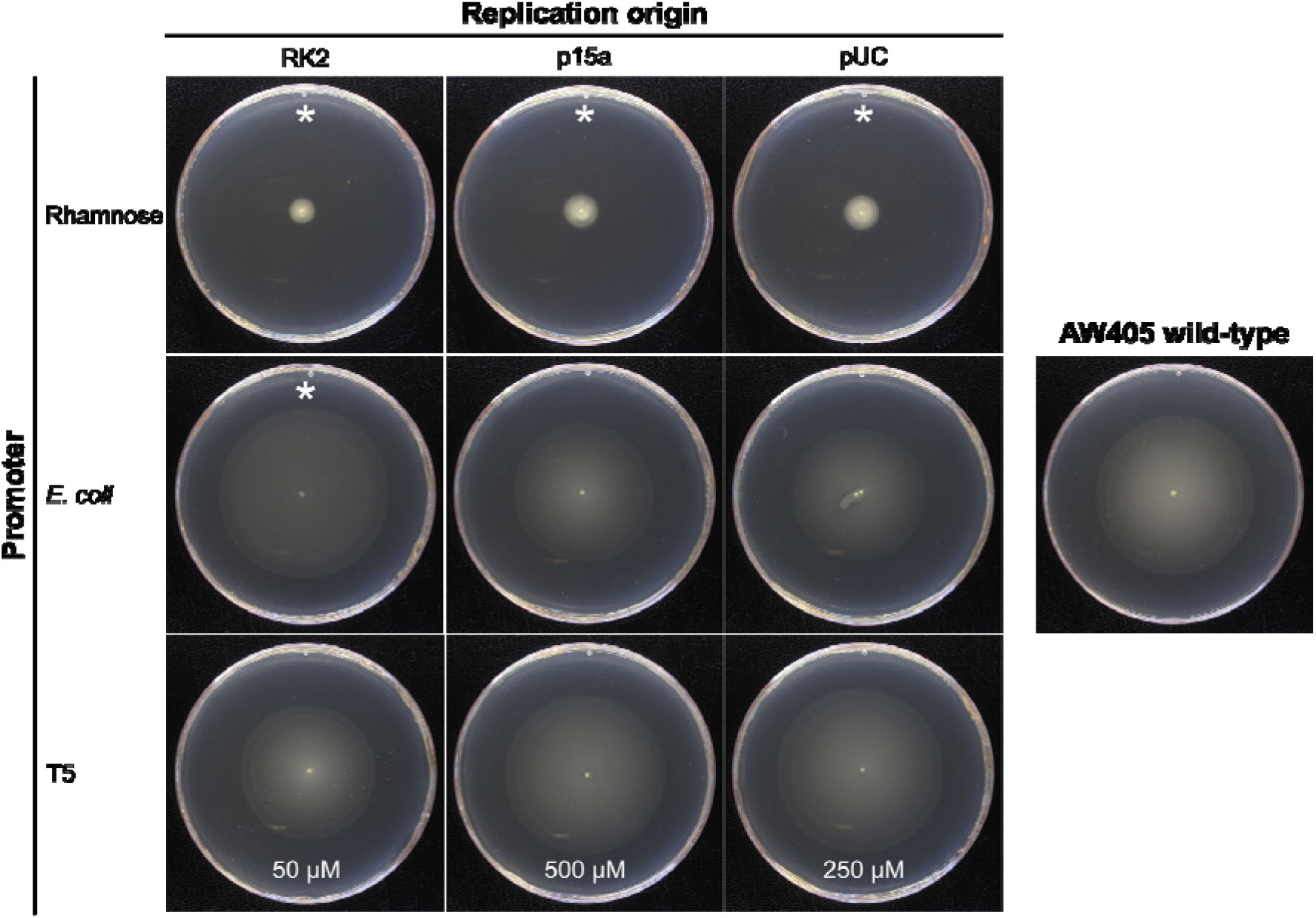
Representative motility agar plates with the conditions that most closely restored wild-type motility for each tested plasmid. All plates were incubated at 30 °C for 24 h. The results with the rhamnose promoter were all achieved with 0.5% rhamnose. Concentrations of IPTG that gave motility closest to wild-type are given at the bottom of the corresponding plates. * = mean motility under these conditions was significantly different to wild-type (T-test; p < 10^−4^)

The *E. coli fliC* promoter is controlled by the endogenous flagellar expression cascade [17] and so is not regulatable. The high copy number plasmid gave the lowest average motility, followed by the medium copy number and then the low copy number (Figure 1 (C)). The high copy number (5.9% decrease) and medium copy number (9.2% increase) plasmids replicated wild-type motility (T-test; p > 0.1; Figure 2). Motility from the low copy number plasmid showed a significant increase of 27.3% over the control (T-test; p < 10^−4^).

### 3.2. Flagellin production

RNA was recovered from triplicate cultures of each strain, including the wild-type and AW405Δ*fliC* controls. AW405Δ*fliC* did not show amplification of a *fliC* target fragment but had similar levels of the endogenous control. Triplicate technical repeats of each reaction showed very low variance. However, significant variation existed between biological repeats, as reflected in the error bars in Figure 3. This might indicate that *fliC* transcription varies widely between growth phases, so that small differences in the OD_600_ at harvest reflected populations of cells on either side of a switch to lower motility in stationary phase.

**Figure 3:**
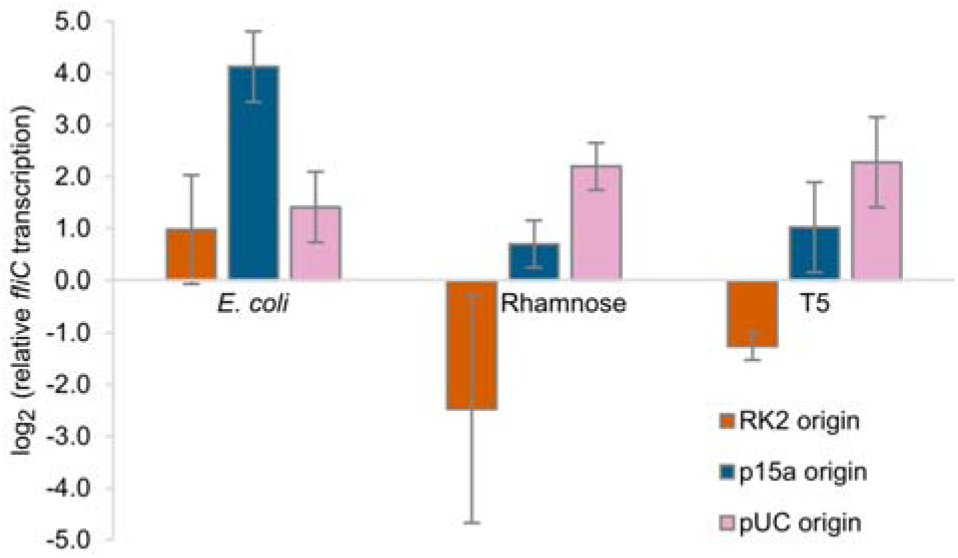
*Log-transformed FliC transcription levels at late-exponential phase relative to wild-type AW405, as measured by RT-qPCR. Wild-type transcription = 0. A* fliC *knockout strain did not give amplification of the target* fliC *fragment. Results are the means of 3 biological replicates each with 3 technical replicates* ± *95% confidence intervals*.

Despite the large variability, a general trend can be seen towards greater transcription as the plasmid copy number increases (Figure 3). Furthermore, the low copy number plasmids showed transcription levels most like the wild-type. The *E. coli fliC* promoter did not show the same trend. Although expression from the rhamnose promoter led to reduced motility, the level of *fliC* transcription was not notably different from the other promoters, although the combination of rhamnose promoter and RK2 replication origin was the only one to give lower transcription levels than the wild-type. Overall, flagellin expression from a plasmid leads to higher transcription levels but only to equal or reduced motility, suggesting that not all the produced flagellin is incorporated into flagella.

### 3.3. Quantification of filament-associated flagellin

Only flagellin that is correctly assembled into a flagellar filament can contribute towards cell motility. Therefore, in addition to quantifying the total transcription of *fliC*, we also recovered the filament-associated protein by heat depolymerisation and compared the concentrations between each strain by densitometry of SDS-PAGE bands.

Interestingly, there was greater correlation between the detected amount of flagellin in filaments and cell motility than there was for transcription. Plasmids utilising the *E. coli* and T5 promoters incorporated similar amounts of flagellin to the wild-type and had similar motilities (Figure 4). When the rhamnose promoter controlled expression, significantly less flagellin was incorporated (T-test; p < 0.001) and motility was also lower. The only other culture to show a significantly different concentration of filament-associated flagellin was the combination of the strong T5 promoter and pUC replication origin, which also decreased (T-test; p < 0.05). Therefore, increased transcription of *fliC* does not necessarily translate to larger quantities of flagellin incorporated into flagellar filaments and may in fact lead to less filament-associated flagellin.

**Figure 4:**
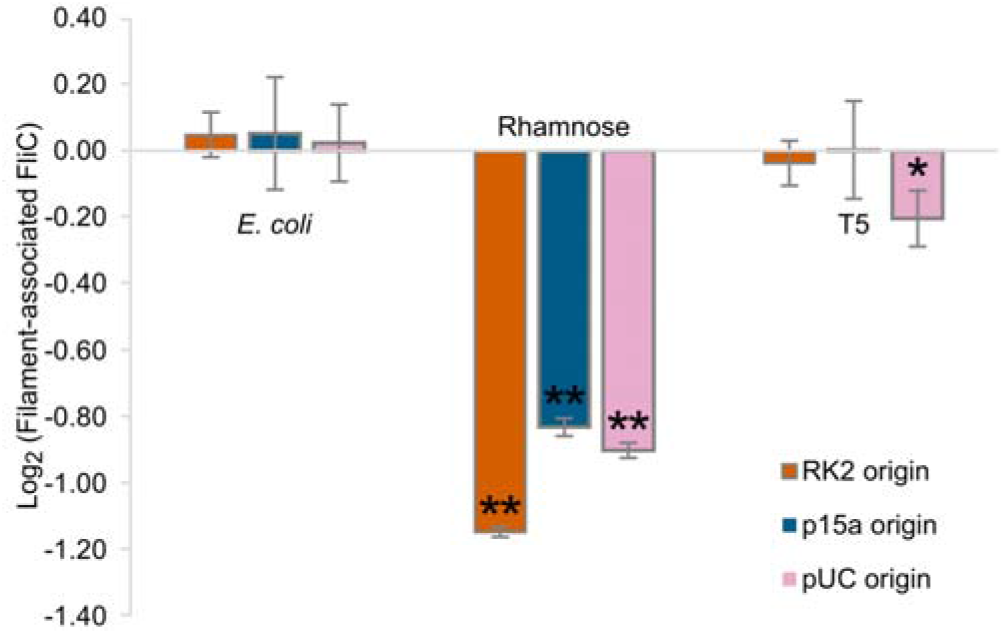
*Log-transformed levels of filament-associated flagellin in each strain tested at late-exponential phase, relative to wild-type AW405. Flagella were depolymerised by heat treatment and the monomers were separated from cells and analysed by densitometry of polyacrylamide gels. Wild-type concentration = 0. A* fliC *knockout strain gave a small signal by densitometry that was not distinguishable from background noise and other proteins, and so is not shown here. Results are the means of 3 biological replicates* ± *SEM*. * = *p* < *0.05;* ** = *p* < *0.001*.

## 4. Discussion

We compared the levels of *E. coli* flagellin expression, incorporation into flagellar filaments, and consequent motility in isogenic AW405Δ*fliC* strains containing a series of nine plasmids with different combinations of promoters and plasmid replication origins. Our aim was to faithfully reinstate wild-type motility to the *fliC* knockout mutant by tuning the expression of wild-type flagellin from the plasmids. The results presented here demonstrate that for a complex system such as the *E. coli* flagellum it can be difficult to predict the effects of manipulating the expression of a single component. Flagellin production and incorporation into flagellar filaments is the final part of a complex gene regulation cascade, involving more than 40 different structural proteins and transcription factors [18]. Since the multiple previous stages in the construction of the flagellum largely determine the magnitude of motility, it is expected that there will not be a simple, direct correlation between flagellin expression and motility.

Due to dosage effects, the low copy number plasmid should most faithfully reproduce the wild-type phenotype as it is most similar to the few copies of *fliC* that are likely to be present on the chromosome (note that during exponential growth, *E. coli* replicates the chromosome faster than cell division and so maintains multiple, rather than single, copies of most genes [19]). Our results showed that plasmid copy number had only a small effect in this regard, indicating that the choice of chassis plasmid for gene complementation is relatively unimportant. Surprisingly, our results indicate that choosing a medium- or high copy number plasmid is slightly preferable to a low copy number when expression is driven by the *fliC* promoter. Our qRT-PCR results suggest that the flagellar regulatory machinery imposes itself when a high copy number plasmid is used, thus limiting flagellin production and allowing normal motility.

The rhamnose and T5 promoters break the connection between *fliC* and the rest of the flagellar structural and regulatory components. With these promoters it is possible to tune cell motility based on flagellin production, from preventing motility in the absence of inducer to replicating wild-type motility with ~250 μM IPTG. Increasing plasmid copy number with the inducible promoters increased *fliC* transcription but did not increase the amount of filament-associated flagellin at the maximum induction strength. It would therefore appear that increasing flagellin production fails to increase motility, either because longer filaments are incapable of providing extra propulsion or because the extra subunits cannot be transported fast enough to assemble. Although inducer concentration was the dominant factor in determining motility, within each concentration bracket there was a small additional effect of plasmid copy number. The low motilities and low concentrations of filament-associated flagellin observed from the rhamnose promoter suggested that it is a relatively weak promoter. This is supported by the observation that higher copy number plasmids allowed slight increases in motility with this promoter.

The choice of promoter and induction strength is crucial. While the rhamnose promoter allowed tight control of motility and was finely tuneable, filament-associated flagellin and motility remained significantly below wild-type under all tested conditions. On the other hand, both the natural *E. coli fliC* promoter and the IPTG-inducible T5 promoter were able to completely rescue the motility phenotype. From this, we would recommend use of the natural promoter for the gene of interest wherever possible because it will be responsive to endogenous control mechanisms. However, if manual control of timing or levels of expression are required, titration of induction strength from an inducible promoter such as T5 may be preferable.

The set of 9 expression plasmids constructed for this study may be useful for future similar studies with other proteins. The plasmids can easily be modified to make use of different promoters by cloning alternative sequences between the SalI and NcoI restriction sites. Any gene of interest can also replace the *fliC* sequence by in-frame insertion between the NcoI and EcoRI restriction sites.

The most predictable way to reinstate a wild-type phenotype may be to re-insert the gene into the chromosome, perhaps under control of an inducible promoter if control over gene expression would be useful. However, chromosomal manipulations are time-consuming, difficult to perform in high-throughput, often leave scar sequences from removal of a selection marker, and generally require the production of a plasmid containing the sequence to be inserted. Therefore, particularly for screening large numbers of genes or gene variants, it is far more convenient to express the desired gene from a plasmid in the first instance and to follow up with more detailed studies of chromosome-based expression if necessary.

This study has provided useful guiding principles for the specific case of restoring motility in *E. coli*. We expect that this will facilitate further investigations into the structure and function of the flagellum and high-throughput biotechnological applications such as the development of flagellar display technology for large, globular protein domains. It should also serve as a reminder that simple optimisation and verification of protein expression strategies should be performed to separate the effects of the protein itself from convoluting factors from the expression system.

## Acknowledgements

Funding: This work was supported by the Quadram Institute Bioscience Biotechnology and Biological Sciences Research Council (https://bbsrc.ukri.org/) Strategic Programme Grant: Microbes in the Food Chain (project number BB/R012504/1).

